# Complementary δ2-protocadherin expression delineates parallel basal ganglia circuits in primates

**DOI:** 10.64898/2026.05.14.725043

**Authors:** Naosuke Hoshina, Miyuki Hoshina, Tadashi Yamamoto, Masahiko Takada

**Affiliations:** Waseda Institute for Advanced Study, Waseda University, Tokyo 162-8480, Japan; Center for the Evolutionary Origins of Human Behavior, Kyoto University, Inuyama, Aichi 484-8506, Japan; Cell Signal Unit, Okinawa Institute of Science and Technology Graduate University, Onna-son, Okinawa 904-0495, Japan

**Keywords:** basal ganglia, striatum, pallidum, substantia nigra, parallel circuits, protocadherin

## Abstract

The basal ganglia (BG) form anatomically and functionally segregated yet integrative parallel circuits, but the molecular mechanisms specifying them remain unclear. We immunohistochemically mapped the expression of three δ2-protocadherin (δ2-PCDH) cell adhesion molecules—PCDH10, PCDH17, and PCDH19—in the BG of macaques. Within the striatum, each PCDH exhibited regional gradients of expression along the rostro–caudal and ventromedial–dorsolateral axes. The three PCDHs showed complementary distributions that continuously delineated molecular boundaries corresponding to functional subdivisions in a graded fashion. Such complementary distributions were also observed in the BG output nuclei. Given that neurons expressing the same δ2-PCDH in distinct BG structures preferentially connect with each other, the three δ2-PCDH expression patterns could define functional territories within parallel BG circuits. Together, the complementary expression of PCDH10, PCDH17, and PCDH19 broadly align with the distinct BG circuits, respectively, suggesting molecular codes underlying the segregated yet integrative parallel organization of the primate BG.

## INTRODUCTION

The basal ganglia (BG) are a collection of subcortical nuclei that play essential roles in regulating a wide range of functions, including motor control, habit formation, reinforcement learning, motivation, cognition, and emotion.^1–3^ This system comprises the striatum (caudate nucleus [Cd], putamen [Pu], and nucleus accumbens [NA]), the pallidum (external [GPe] and internal [GPi] segments of the globus pallidus, and ventral pallidum [VP]), the subthalamic nucleus (STN), and the substantia nigra (pars compacta [SNc] and pars reticulata [SNr]). The striatum receives excitatory input from the cerebral cortex and relays information through two principal, direct and indirect, pathways.^4–6^ In the former pathway, striatal projection neurons innervate the GPi and SNr, facilitating motor output. In the latter pathway, on the other hand, striatal neurons project to the GPe, which then connects to the GPi and SNr via the STN, resulting in net inhibition of output activity. Both pathways converge onto the thalamus, forming cortico–BG–thalamocortical loops that are crucial for achieving multiple functions. Dysfunctions of these circuits have been implicated in a variety of neurological and psychiatric disorders, including Parkinson’s disease, Huntington’s disease, autism spectrum disorder (ASD), obsessive–compulsive disorder (OCD), schizophrenia, and depression.^2,7–12^

The cortico–BG networks are organized into parallel circuits with a topographic arrangement of functionally independent domains, reflecting the classical view of anatomical and functional segregation.^13,14^ More recent studies extend this model, suggesting that these circuits are not strictly segregated but form continuous parallel networks with integrative interactions, including overlapping connectivity and functional crosstalk across territories.^15–19^ In primates, these parallel cortico–BG networks are classically divided into three circuits^20,21^: the sensorimotor circuit, which projects from sensorimotor cortical areas to the caudal striatum and mediates sensorimotor functions; the associative circuit, which projects from prefrontal cortical areas to the rostral striatum and supports executive and cognitive functions; and the limbic circuit, which projects from limbic cortical areas to the ventral striatum and governs emotional and motivational processes. Importantly, recent electrophysiological studies have demonstrated that within this framework of parallel information processing, flexible associative circuits favor cognitive decision-making, whereas stable sensorimotor circuits mediate habitual actions.^22,23^ A comparable, although species-specific, topographic organization is also observed in rodents.^24–26^

Despite this well-characterized anatomical and functional framework, it remains unclear whether these parallel BG circuits are genetically represented, and which molecular codes define them. Molecular neuroanatomical approaches are essential for dissecting the brain at high resolution, because region- and cell-type–specific gene expression often reflects the underlying circuit architecture.^27^ For instance, in the direct and indirect pathways, D1 and D2 dopamine receptors are specifically expressed in their respective striatal medium spiny projection neurons, acting as central mediators of opponent circuit regulation.^28,29^ In addition, the μ-opioid receptor shows striosome-specific expression, while calbindin marks matrix compartments, together defining striatal striosome/matrix domains with distinct connections and functions.^30,31^ These largely exclusive expression patterns enable clear delineation of pathways and compartments in the BG, but boundaries between the parallel BG circuits are often continuous, making molecular specification challenging. In rodents, members of the cadherin family exhibit region-specific gradients of expression in the BG,^32,33^ suggesting a potential role in specifying the parallel BG circuits. Notably, our studies identified δ2-protocadherin (δ2-PCDH) family members PCDH10 and PCDH17 as key molecules, exhibiting complementary expression and selective neuron–neuron interactions in mice, supporting their roles in establishing parallel BG networks.^33^ However, it remains unknown whether δ2-PCDH members similarly specify parallel BG circuits in primates.

Our primary objective is to elucidate the organizational mechanisms and the molecular events that underly the anatomical segregation of parallel BG circuits. The present study addresses this issue by mapping the expression of the three δ2-PCDH proteins—PCDH10, PCDH17, and PCDH19—in the BG of infant macaques. The use of infant monkeys is crucial because the expression of these d2-PCDHs peaks and undergoes dynamic changes during early development, when synaptogenesis is most active; therefore, using infant subjects—who are at the peak of this critical molecular transition—is essential, in contrast to adults in whom expression is typically reduced. This approach is consistent with our previous work examining PCDH17 expression in infant primates^33^, ensuring the comparability of findings across studies. Using immunohistochemistry on serial sections, we observed that these δ2-PCDHs display partial overlaps yet complementary distributions within the striatum, largely consistent with functional subdivisions of the limbic, associative, and sensorimotor territories. Such complementary patterns were also observed in the pallidum and substantia nigra, suggesting that individual neurons expressing the same δ2-PCDH form each parallel network of the BG. These findings provide new insights into the molecular delineation of functional architecture in the primate BG, indicating that δ2-PCDHs contribute to the specification and organization of the parallel BG circuits.

## RESULTS

### δ2-Protocadherin Members: PCDH10, PCDH17, and PCDH19

The primate BG are organized into parallel circuits that process limbic, associative, and sensorimotor information, exhibiting a continuous topographic arrangement within the striatum (Figure 1).^12,16,17,19^ This organization comprises three major circuits: the limbic circuit located in the ventral striatum, the associative circuit positioned in the rostral striatum, and the sensorimotor circuit extending across the caudal striatum. Each of these territories forms continuous gradients that partially overlap with one another, creating transitional zones that facilitate functional integration and communication between the parallel circuits.^15,16,18^

**Figure 1.**
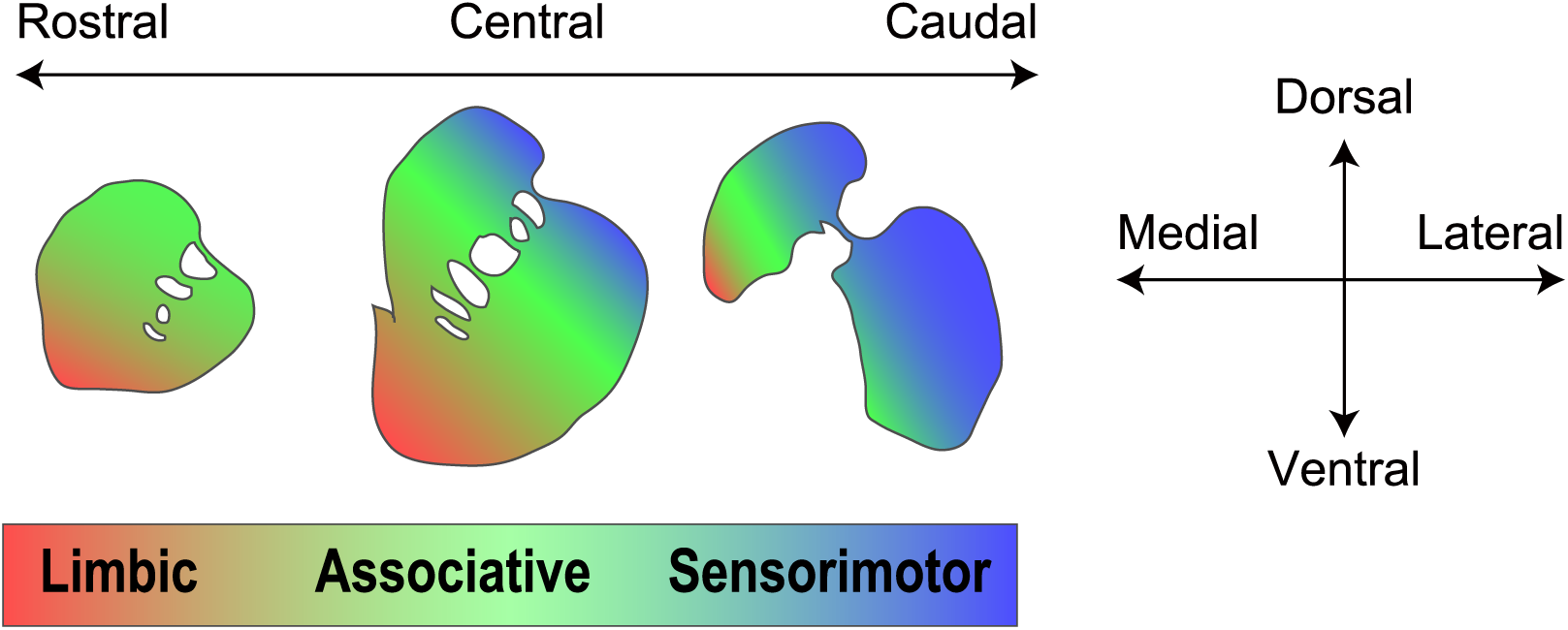
Organization of parallel BG circuits in the primate striatum. Schematic diagram illustrating the functional territories in parallel circuits of the primate striatum, showing the continuous topographic organization of the limbic (ventral striatum; red), associative (rostral striatum; green), and sensorimotor (caudal striatum; blue) subdivisions. Color gradients indicate partial overlap and potential integrative interactions between these circuits. The schematic was adapted from previous studies.^12,16^

To investigate the molecular architecture underlying this parallel circuit organization, we focused on non-clustered δ2-PCDHs, specifically PCDH10, PCDH17, and PCDH19. Unlike the clustered α-, β-, and γ-PCDHs, which are encoded by large gene clusters that generate multiple extracellular cadherin domains, non-clustered PCDHs are encoded by individual genes, each producing a single extracellular domain^34^. This fundamental genomic distinction is accompanied by clear differences in gene expression patterns, molecular properties, and physiological functions.^34,45^ Non-clustered PCDHs are single-pass transmembrane cell adhesion molecules broadly expressed throughout the central nervous system and are classified into two subfamilies: δ1 (PCDH1, PCDH7, PCDH9, PCDH11, PCDH20) and δ2 (PCDH8, PCDH10, PCDH12, PCDH17, PCDH18, PCDH19) (Figure 2A).^34^ Sequence alignment revealed that PCDH10, PCDH17, and PCDH19 are highly conserved across mouse (*Mus musculus*), monkey (*Macaca mulatta*), and human (*Homo sapiens*), representing the most closely related members within the δ2-PCDH subfamily. All three δ2-PCDHs share six extracellular cadherin repeats (EC domains) and two conserved cytoplasmic motifs (CM1 and CM2) (Figure 2B). Functionally, these δ2-PCDHs mediate homophilic trans-interactions, in which identical PCDH molecules bind to each other through EC1–EC4 (e.g., PCDH10–PCDH10), but they do not engage in heterophilic interactions with other δ2-PCDH members (e.g., PCDH10–PCDH17).^33,36^ Within their cytoplasmic domains, all three δ2-PCDHs are associated with the WAVE regulatory complex, suggesting that they regulate largely overlapping intracellular signaling pathways.^35,37^ Phenotypic analyses using *Pcdh10*, *Pcdh17*, and *Pcdh19* knockout mice indicate that, rather than controlling axonal projections, these molecules primarily regulate the development, function, and plasticity of synapses within specific neural circuits.^33,38,39^ Collectively, these conserved structural and functional features suggest that PCDH10, PCDH17, and PCDH19 play critical roles in selective neuron–neuron recognition and in the precise molecular assembly of BG circuits.

**Figure 2.**
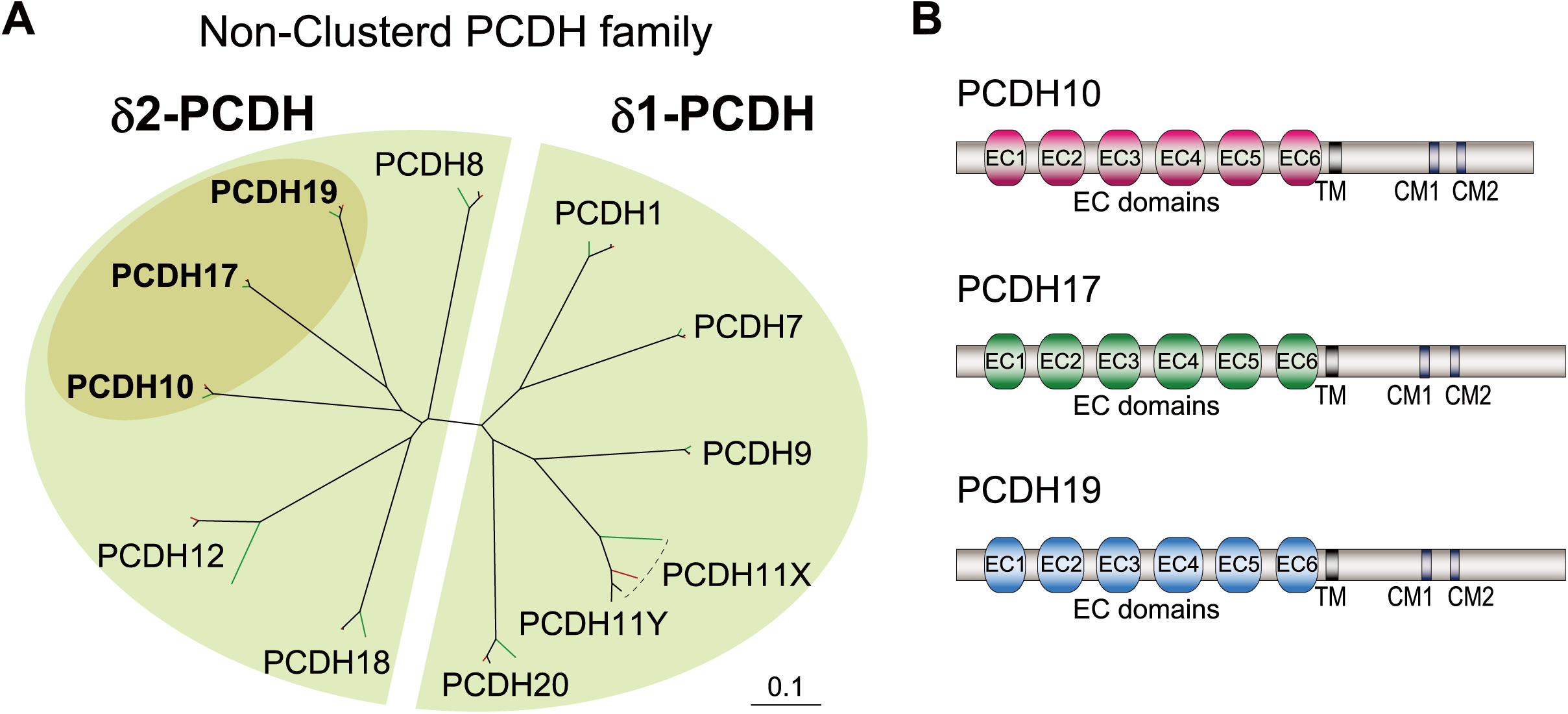
PCDH10, PCDH17, and PCDH19 belong to the δ2-protocadherin family. (A) Phylogenetic tree of non-clustered protocadherins (PCDHs) from mouse (Mus musculus; green), monkey (Macaca mulatta; red), and human (Homo sapiens; black), including δ1- and δ2-PCDH family members, based on full-length protein sequences. Multiple sequence alignment and phylogenetic analysis were performed using Clustal Omega, and the resulting tree was visualized with TreeView. The scale bar indicates amino acid substitution rate per ten residues. (B) Schematic representation of the protein structures of monkey PCDH10 (XP_002804241.2; isoform X2), PCDH17 (NP_001252640.1), and PCDH19 (NP_001252617.1). EC, extracellular cadherin-like domain; TM, transmembrane domain; CM1/2, conserved motifs among δ-PCDHs.

### Complementary Expression of PCDH10, PCDH17, and PCDH19 in the Striatum

To determine whether the δ2-PCDH members PCDH10, PCDH17, and PCDH19 delineate molecularly discrete regions corresponding to functional territories within the BG, we performed immunohistochemical analyses on serial coronal sections from rhesus monkeys. Representative immunostaining images of the striatum, including the caudate nucleus (Cd), putamen (Pu), and nucleus accumbens (NA), are presented in Figure 3 (see the striatal regions in Figure 3A). The corresponding color-coded composite maps in Figure 4 illustrate the overall spatial distribution of PCDH10 (red), PCDH17 (green), and PCDH19 (blue), as well as their pairwise and triple-color merged patterns. The three δ2-PCDHs exhibited distinct and continuous topographic expression gradients along both the rostro–caudal and dorsolateral–ventromedial axes.

**Figure 3.**
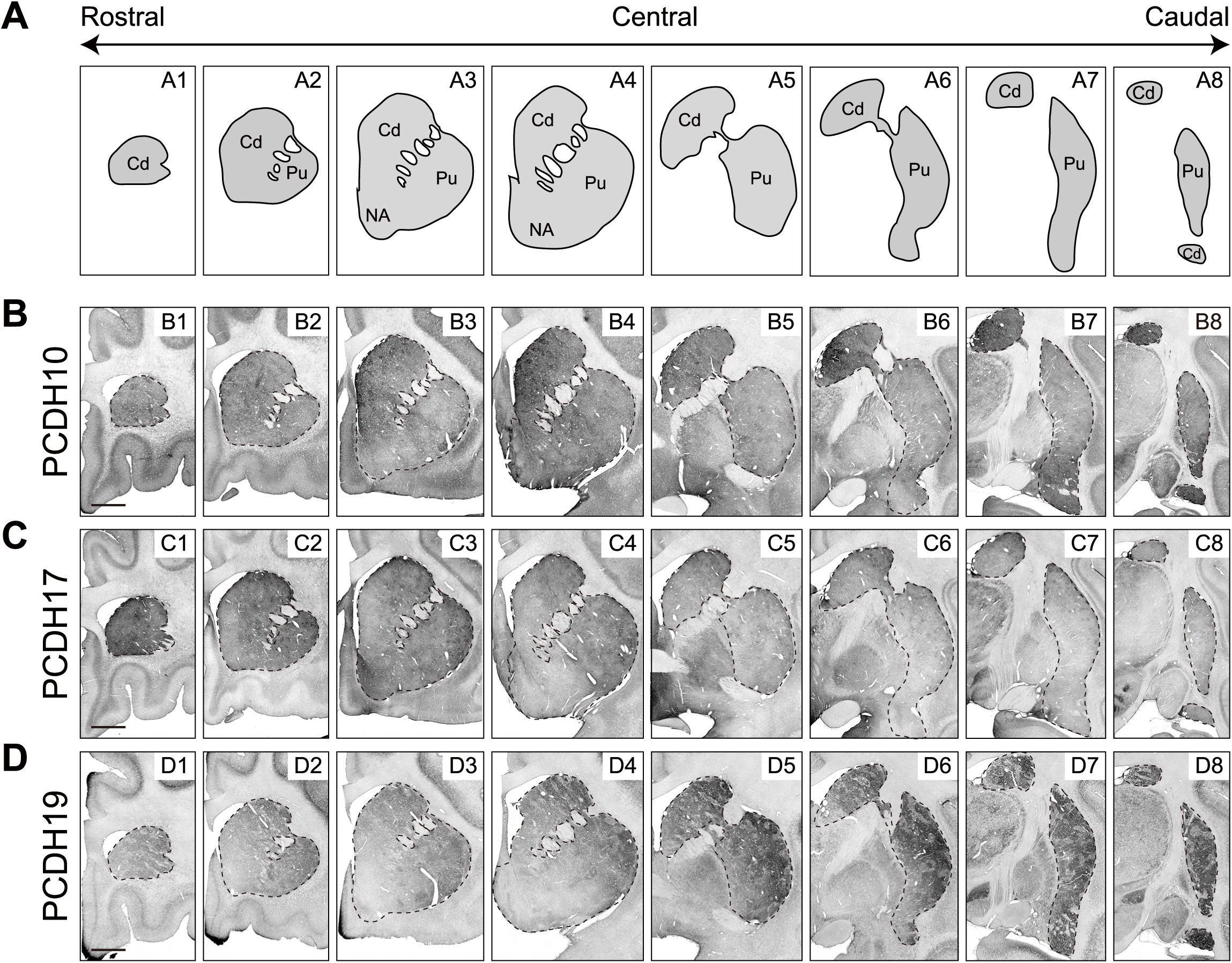
Distribution of PCDH10, PCDH17, and PCDH19 in the monkey striatum. (A) Schematic of coronal sections through the striatum, including the caudate nucleus (Cd), putamen (Pu), and nucleus accumbens (NA), along the rostral–central–caudal axis. (B–D) Immunostaining for PCDH10, PCDH17, and PCDH19 in serial coronal sections of the striatum from a 1- to 2-month-old rhesus monkey. Each δ2-PCDH shows distinct rostral–caudal and dorsolateral–ventromedial gradients: (B) PCDH10 is highest in the ventromedial striatum, including the NA; (C) PCDH17 is highest in the rostral striatum; (D) PCDH19 is highest in the caudal striatum. Partial overlaps create continuous gradients between territories. Scale bar, 3 mm (B–D).

**Figure 4.**
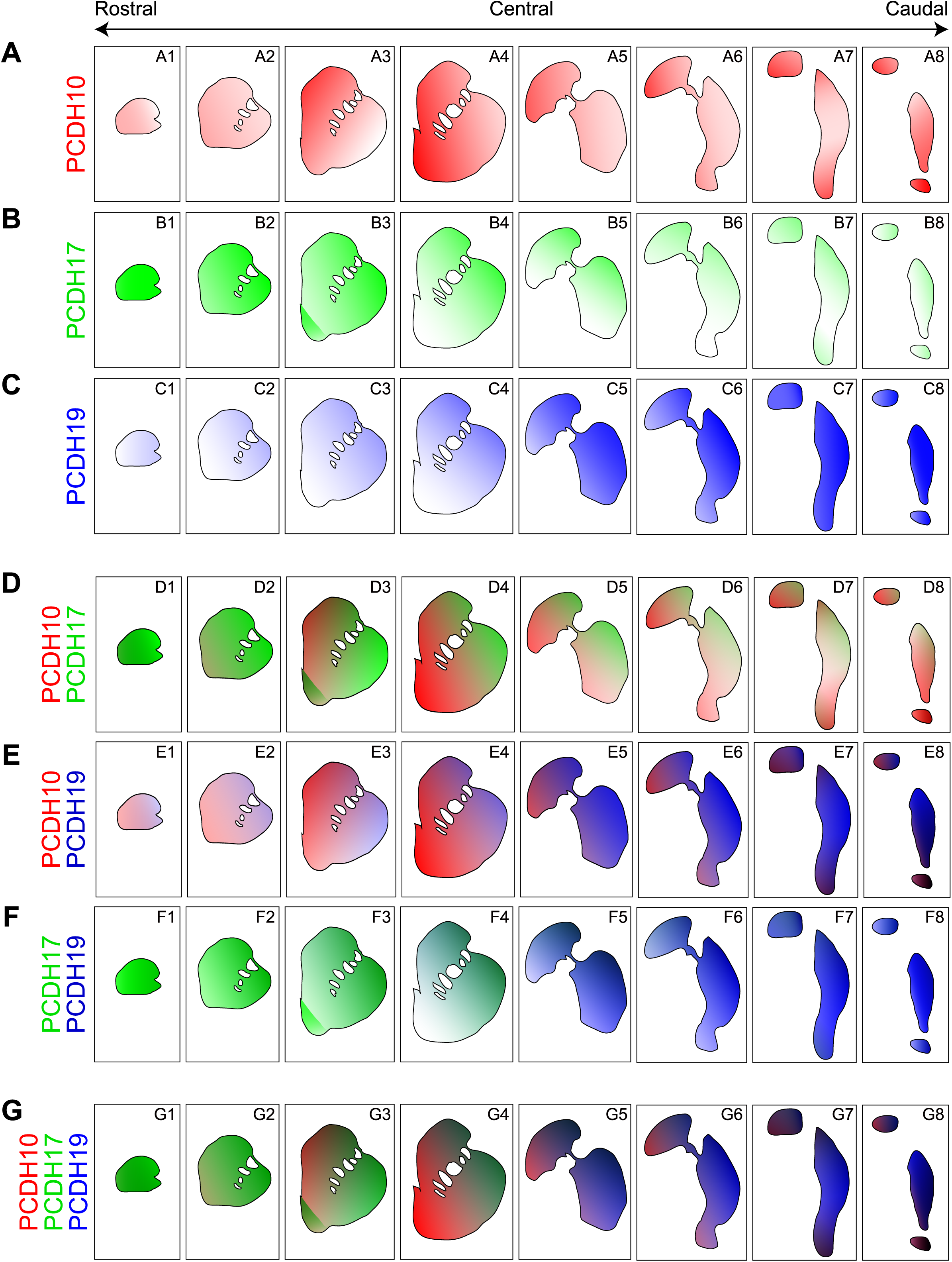
Complementary expression of PCDH10, PCDH17, and PCDH19 in the striatum. Color-coded maps (PCDH10: red; PCDH17: green; PCDH19: blue) based on the immunostaining shown in Figure 3. (A–C) Single-color maps for each δ2-PCDH show regional predominance and spatial gradients: (A) PCDH10 is enriched in the ventromedial striatum, gradually decreasing toward the dorsolateral striatum; (B) PCDH17 is concentrated in the rostral striatum, with gradients along both rostral–caudal and dorsolateral–ventromedial axes; (C) PCDH19 is highest in the caudal striatum, gradually decreasing along both caudal–rostral and dorsolateral–ventromedial axes. (D–F) Dual-color merged maps illustrate pairwise complementarity: (D) PCDH10/PCDH17, ventromedial vs. dorsolateral in the rostral–central striatum; (E) PCDH10/PCDH19, ventromedial vs. dorsolateral in the central–caudal striatum; (F) PCDH17/PCDH19, rostral vs. caudal complementarity. (G) Triple-color merged map (PCDH10/PCDH17/PCDH19) reveals largely complementary distributions across striatal subregions, with gradual transitions and partial overlaps in intermediate regions.

PCDH10 expression was the highest in the ventromedial portion of the central striatum, including the NA and medial caudate putamen (CPu) (Figures 3B4 and 4A4). Strong signals extended into the caudal Cd (Figures 3B5–8 and 4A5–8). Overall, the signal intensity gradually decreased toward the dorsolateral striatum, forming a smooth ventromedial–dorsolateral gradient that characterizes the ventral subdivision of the striatum (Figures 3B and 4A). PCDH17 showed strong expression in the rostral striatum, specifically in the head region (the most anterior tip of the caudate and putamen) (Figures 3C1 and 4B1). Its intensity gradually decreased toward more caudal regions (Figures 3C and 4B). A prominent dorsolateral–ventromedial gradient was evident in the CPu, most pronounced in the central region (Figures 3C4 and 4B4). Additionally, a secondary region of relatively strong expression was detected in the rostral portion of the NA (Figures 3C3 and 4B3), where PCDH10 expression appeared comparatively weaker (Figures 3B3 and 4A3). PCDH19 exhibited the highest expression in the caudal CPu, including the caudal tail region (the most posterior part of the caudate) (Figures 3D8 and 4C8). Its expression gradually decreased toward rostral regions (Figures 3D and 4C), demonstrating a reciprocal rostro–caudal gradient to that of PCDH17. A dorsolateral–ventromedial gradient was also observed in the CPu (Figures 3D2–4 and 4C2–4), which was similar to the pattern observed for PCDH17.

The pairwise merged maps (Figure 4D–F) further demonstrate the broadly complementary spatial organization among these δ2-PCDH expression territories. Overlaps between adjacent expression zones were generally confined to transitional regions, indicating gradual molecular transitions rather than abrupt boundaries, thereby supporting the model of continuous gradients. Specifically, the PCDH10/PCDH17 map reveals an organization characterized by a ventromedial–dorsolateral transition in the central striatum (Figure 4D). Within the NA, PCDH17 exhibits high expression in the rostral part, whereas PCDH10 shows high expression in the caudal part, clearly illustrating complementary distributions within this limbic region (Figure 4D3,4). Notably, both PCDH10 and PCDH17 show relatively reduced expression in the caudal Pu (Figure 4D5–8). Conversely, the PCDH10/PCDH19 map highlights a ventromedial–dorsolateral transition connecting the two territories (Figure 4E), especially central CPu (Figure 4E4). A key feature of this map is the relatively low expression of both PCDH10 and PCDH19 in the rostral regions (Figure 4E1,2). The PCDH17/PCDH19 map illustrates a rostral–caudal transition (Figure 4F), where a significant overlap is observed along the dorsolateral CPu in the central region (Figure 4F4), effectively bridging the rostral PCDH17-dominant and caudal PCDH19-dominant territories. The triple-color merged patterns reveal continuous and complementary δ2-PCDH expression gradients across the striatum, with overlap in intermediate regions (Figure 4G). These findings indicate that δ2-PCDHs establish distinct molecular territories that likely correspond to the ventral (limbic), rostral (associative), and caudal (sensorimotor) subdivisions of the primate striatum.

### Complementary Expression of PCDH10, PCDH17, and PCDH19 in the Pallidum

We next examined the expression of the δ2-PCDH family within the pallidum, which includes the globus pallidus external segment (GPe), globus pallidus internal segment (GPi), and ventral pallidum (VP) (Figure 5A). Each δ2-PCDH exhibited distinct topographic enrichment patterns across these nuclei (Figures 5–7), showing complex distributions rather than uniform rostro-caudal or ventromedial–dorsolateral gradients.

**Figure 5.**
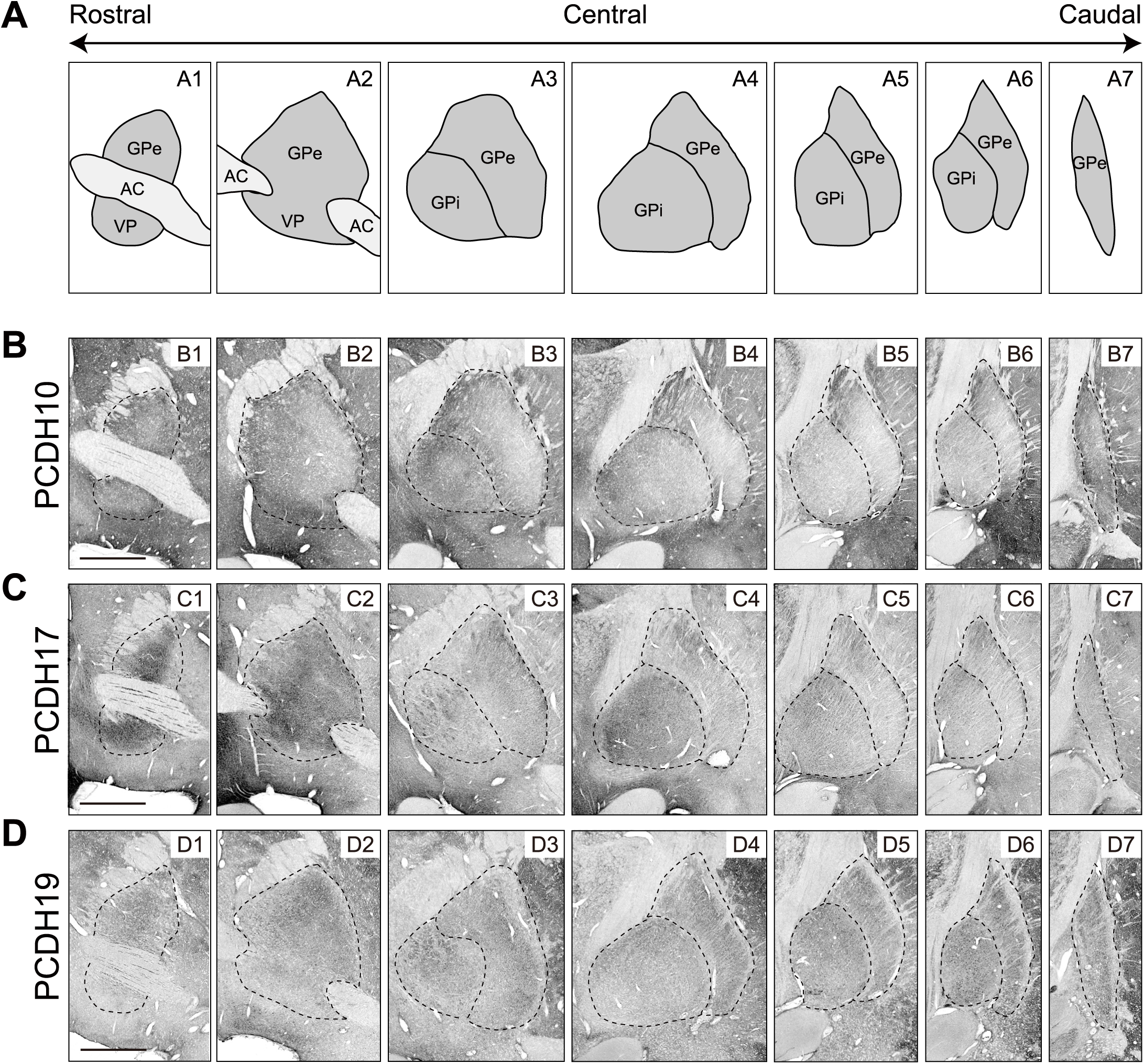
Distribution of PCDH10, PCDH17, and PCDH19 in the monkey pallidum. (A) Schematic of coronal sections through the pallidum, including the external globus pallidus (GPe), internal globus pallidus (GPi), and ventral pallidum (VP), along the rostrocaudal axis. AC, anterior commissure. (B–D) Immunostaining for PCDH10, PCDH17, and PCDH19 in serial coronal sections of the pallidum from a 1- to 2-month-old rhesus monkey. Each δ2-PCDH shows distinct region-specific patterns: (B) PCDH10 is enriched in the peripheral regions of the pallidum; (C) PCDH17 is enriched in the rostral pallidum; (D) PCDH19 is enriched in the caudal pallidum. Partial overlaps create continuous gradients between territories. Scale bar, 3 mm (B–D).

Focusing on the GPe and VP (Figures 5 and 6), the three δ2-PCDHs displayed complementary distributions. PCDH10 was expressed in both GPe and VP, with strong expression in peripheral regions along the entire rostrocaudal axis (Figures 5B and 6A). PCDH17 was also present in both nuclei, enriched in inner regions of the rostral GPe/VP (Figures 5C1,2 and 6B1,2), but shifting toward the dorsolateral edge in caudal GPe (Figures 5C3–7 and 6B3–7). Conversely, PCDH19 was observed primarily in the GPe, showing minimal expression in the VP and enrichment in the inner regions (Figures 5D and 6C), with lower levels rostrally (Figures 5D1 and 6C1). Dual-color merged maps (Figure 6D–F) further illustrate these pairwise complementary patterns. The PCDH10/PCDH17 merge revealed peripheral versus inner complementarity in the rostral GPe/VP (Figure 6D), the PCDH10/PCDH19 merge showed peripheral versus inner complementarity in the caudal GPe (Figure 6E), and the PCDH17/PCDH19 merge highlighted rostral versus caudal complementarity across the GPe/VP (Figure 6F). Finally, the triple-color merged map demonstrated complementary expression of the three δ2-PCDHs throughout the GPe/VP with partial overlaps confined to transitional zones between dominant regions (Figure 6G). PCDH10 occupied peripheral territories, PCDH17 marked rostral inner territories, and PCDH19 predominated in caudal inner territories, revealing their dominant territories exhibited complementary patterns.

**Figure 6.**
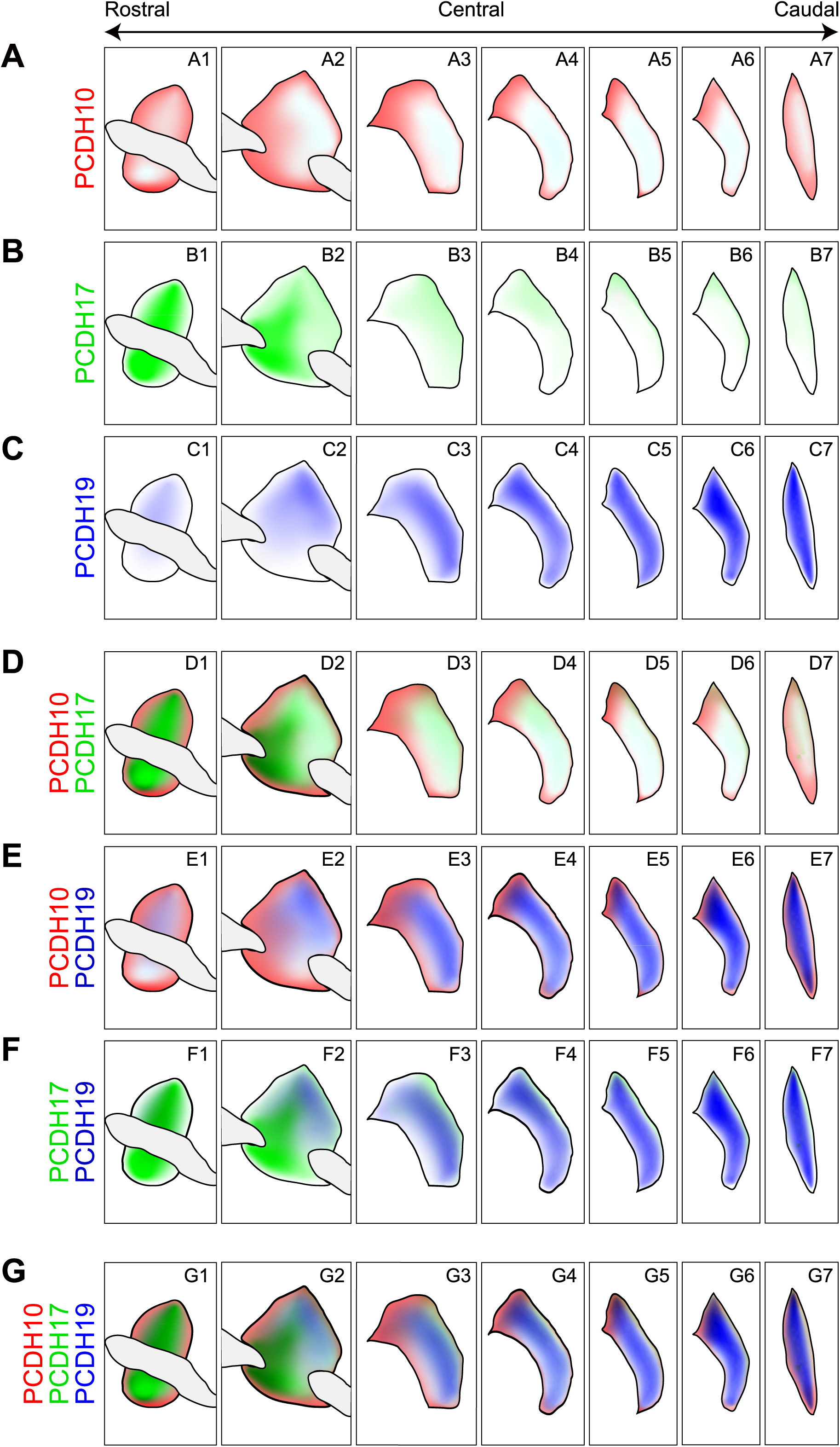
Complementary expression of PCDH10, PCDH17, and PCDH19 in the GPe/VP. Color-coded maps (PCDH10: red; PCDH17: green; PCDH19: blue) based on immunostaining shown in Figure 5. (A–C) Single-color maps for each δ2-PCDH show regional predominance: (A) PCDH10 is strong in the peripheral regions across rostral–caudal GPe/VP; (B) PCDH17 is enriched in the inner regions rostrally in GPe/VP, but at the dorsolateral edge caudally; (C) PCDH19 is enriched in the inner regions of the GPe, with lower levels rostrally. (D–F) Dual-color merged maps illustrate pairwise complementarity: (D) PCDH10/PCDH17 show peripheral vs. inner complementarity in the rostral GPe/VP; (E) PCDH10/PCDH19 show peripheral vs. inner complementarity in the caudal GPe; (F) PCDH17/PCDH19 show rostral vs. caudal complementarity in the GPe/VP. (G) The triple-color merged map (PCDH10/PCDH17/PCDH19) reveals largely complementary expression patterns across the GPe, with partial overlaps in transitional zones.

The GPi exhibited a distinct organizational pattern of the three δ2-PCDHs (Figure 7). PCDH10 was strongly expressed throughout the rostral and central GPi (Figure 7A), with relatively higher intensity in ventromedial domains (Figure 7A2). PCDH17 was also enriched in the rostral and central GPi (Figure 7B1,2), but its expression declined toward caudal sections (Figure 7B3,4). Conversely, PCDH19 was predominantly expressed in the caudal GPi (Figure 7C), showing only low expression rostrally (Figure 7C1). Dual-color merged maps further illustrate these spatial relationships: the PCDH10/PCDH17 merge highlights the ventromedial concentration of PCDH10 compared with the more central/lateral distribution of PCDH17 (Figure 7D). The PCDH10/PCDH19 and PCDH17/PCDH19 merges, in turn, reveal clear rostral–caudal complementarity (Figure 7E,F). Finally, the triple-color map (Figure 7G) demonstrates a largely complementary expression pattern across the GPi, with partial overlaps confined to transitional zones. Overall, these results indicate that δ2-PCDHs define molecularly distinct subregions across all major pallidal nuclei (GPe, GPi, VP), reflecting the topographic organization of parallel striatal outputs, analogous to the patterns observed in the striatum.

**Figure 7.**
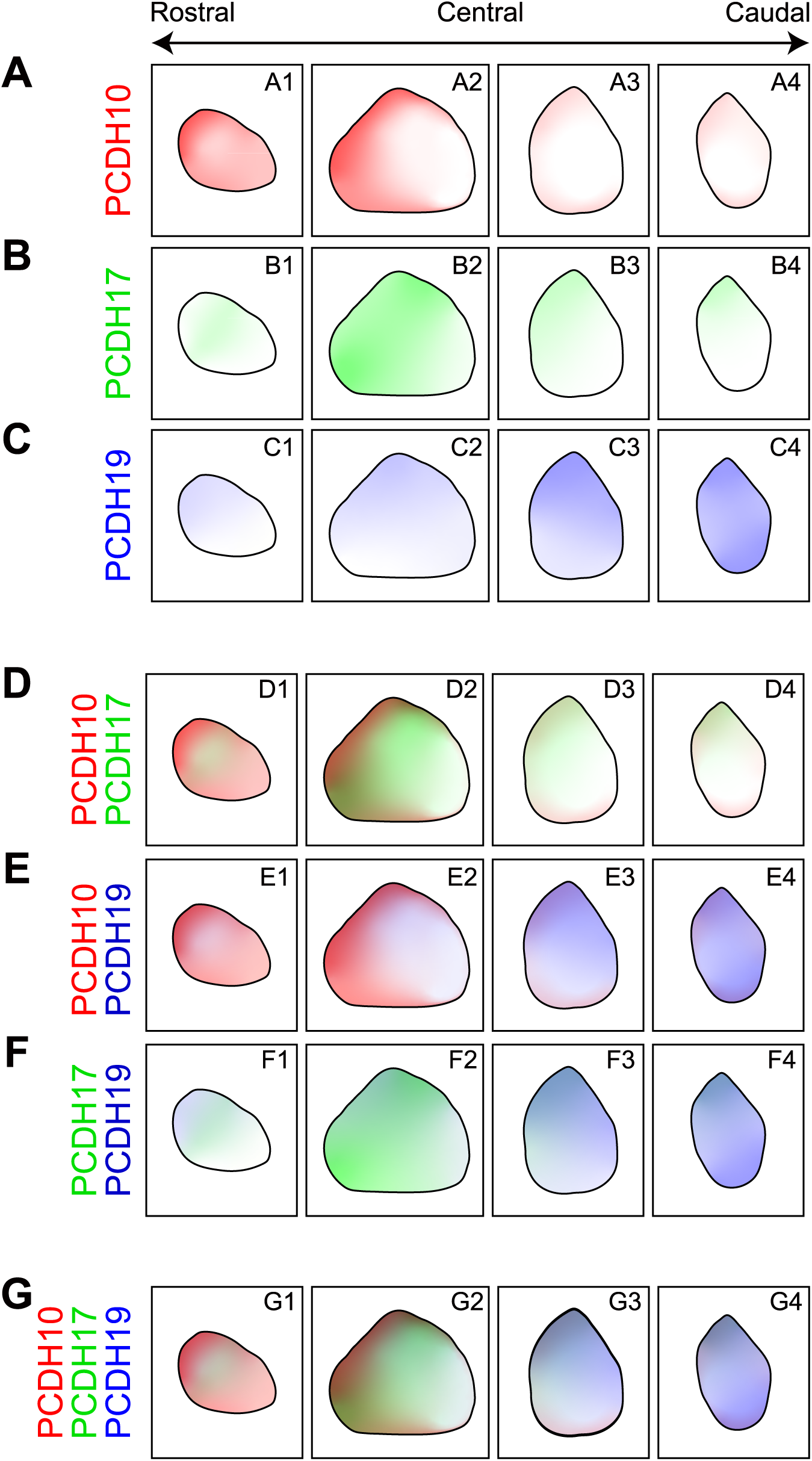
Complementary expression of PCDH10, PCDH17, and PCDH19 in the GPi. Color-coded maps (PCDH10: red; PCDH17: green; PCDH19: blue) based on immunostaining shown in Figure 5B3–B6, C3–C6 and D3–D6. (A–C) Single-color maps: (A) PCDH10 is strongly expressed in the rostral and central regions of the GPi, with relatively higher intensity in ventromedial domains. (B) PCDH17 is strongly expressed in the rostral and central GPi, with intensity declining sharply toward caudal sections. (C) PCDH19 is strongly enriched in the caudal GPi, with low expression in rostral sections.}(D–F) Dual-color merged maps illustrate pairwise complementarity. (D) PCDH10/PCDH17 highlights the ventromedial concentration of PCDH10 compared with the more central/lateral distribution in the rostral–central GPi. (E) PCDH10/PCDH19 clearly demonstrates rostral–caudal complementarity in the GPi. (F) PCDH17/PCDH19 also reveals prominent rostral–caudal complementarity in the GPi. (G) The triple-color merged map (PCDH10/PCDH17/PCDH19) reveals largely complementary expression patterns across the GPi, with partial overlaps confined to transitional zones.

### Complementary Expression of PCDH10, PCDH17, and PCDH19 in the Substantia Nigra Pars Reticulata

To examine whether δ2-PCDH–defined molecular domains extend to BG output nuclei, we analyzed the SNr, the downstream target of the direct pathway (Figures 8 and 9). The SNr lacks the organized, tier-like dopaminergic arrangement characteristic of the neighboring SNc, and this anatomical feature was used as a landmark to distinguish the SNc/SNr boundary. We did not include the STN, SNc, or ventral tegmental area (VTA) in our analysis because our focus is on δ2-PCDH–based projection patterns originating from the striatum, encompassing the striato-pallidal and striato-nigral pathways. Although the STN shows strong PCDH19 expression, the PCDH10/PCDH17 patterns are not sufficient to characterize molecular subdomains. The SNc and VTA, consisting mainly of dopaminergic modulatory neurons, fall outside the scope of the circuitry addressed in this study.

**Figure 8.**
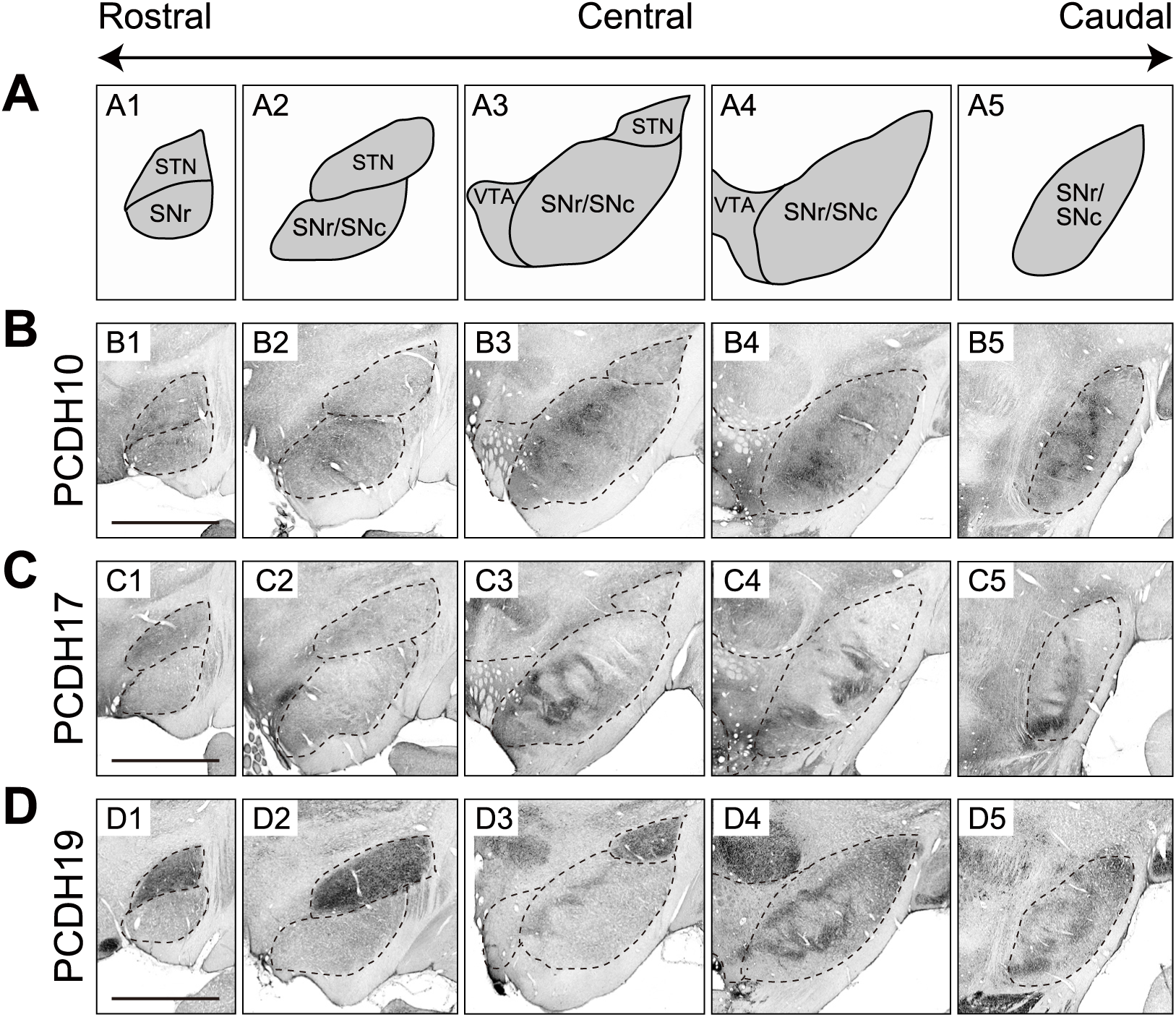
Distribution of PCDH10, PCDH17, and PCDH19 in the monkey substantia nigra pars reticulata (SNr) (A) Schematic coronal sections illustrating the subthalamic nucleus (STN), substantia nigra—pars reticulata (SNr) and pars compacta (SNc)—and ventral tegmental area (VTA) along the rostro-caudal axis. The present study focused on the SNr, which lacks the tier-like organization observed in the dopaminergic neurons of the SNc; neighboring nuclei such as the STN, SNc, and VTA were not analyzed in detail. (B–D) Immunostaining for PCDH10, PCDH17, and PCDH19 in serial coronal sections of the substantia nigra from a 1- to 2-month-old rhesus monkey. Each δ2-PCDH shows a distinct region-specific distribution within the SNr: (B) PCDH10 is preferentially expressed in the medial and dorsal SNr; (C) PCDH17 is preferentially expressed in the ventral SNr; (D) PCDH19 shows highest expression in the caudolateral SNr. Partial overlaps form continuous gradients between territories. Scale bar, 3 mm.

**Figure 9.**
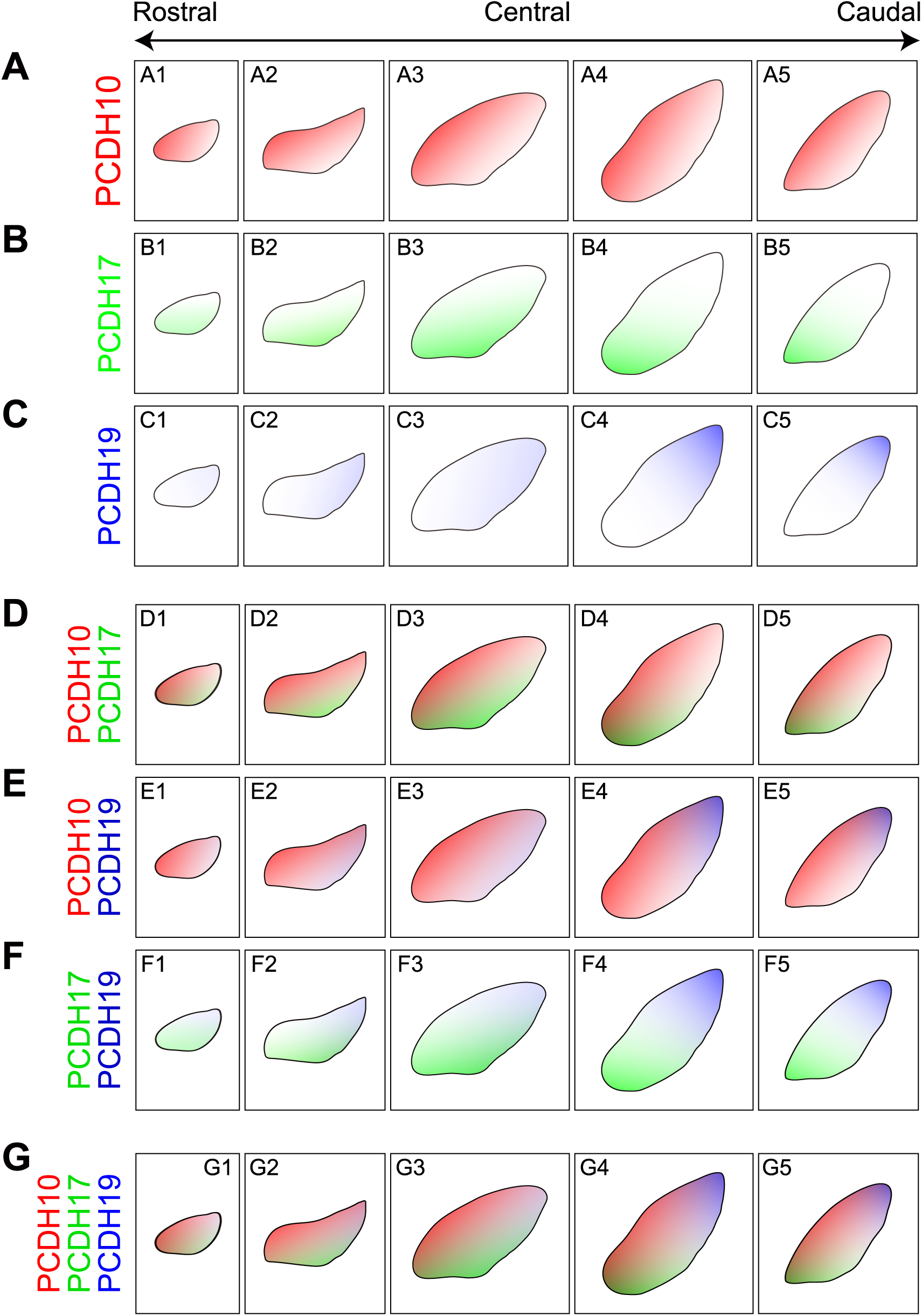
Complementary expression of PCDH10, PCDH17, and PCDH19 in the SNr. Color-coded maps (PCDH10: red; PCDH17: green; PCDH19: blue) are based on immunostaining shown in Figure 8. (A–C) Single-color maps illustrate the regional predominance: (A) PCDH10 is broadly distributed but shows relatively higher expression in the medial and dorsal regions of the SNr. (B) PCDH17 is strongly expressed in the ventral regions of the SNr. (C) PCDH19 is enriched in the caudolateral regions, showing a dorsal bias and low expression rostrally. (D–F) Dual-color merged maps illustrate pairwise complementarity: (D) PCDH10/PCDH17 reveals a dorsal/medial bias of PCDH10 and ventral predominance of PCDH17 in the rostral SNr. (E) PCDH10/PCDH19 demonstrates medial–lateral complementarity in the caudal SNr. (F) PCDH17/PCDH19 highlights ventral–dorsal complementarity. (G) The triple-color merged map (PCDH10/PCDH17/PCDH19) reveals complementary expression patterns across the SNr, with overlaps confined to transitional zones.

Each δ2-PCDH showed a distinct, gradient-like distribution within the SNr (Figures 8B–D and 9A–C). PCDH10 was broadly distributed, with higher expression in medial and dorsal regions of the SNr (Figures 8B and 9A). PCDH17 was concentrated in the ventral SNr along the rostro–caudal axis (Figures 8C and 9B), whereas PCDH19 was most highly expressed in the caudolateral SNr, displaying a strong dorsal bias and minimal rostral expression (Figures 8D and 9C), effectively forming a molecular boundary between dorsal and ventral territories.

Dual-color merged maps revealed dynamic, pairwise complementarity along the rostro–caudal axis (Figure 9D–F). In rostral sections, where PCDH19 expression was low, segregation was primarily driven by PCDH10 and PCDH17, with PCDH10 biased dorsally and medially and PCDH17 predominating ventrally, thereby revealing a subtle multi-axial micro-segregation within the ventral compartment (Figure 9D). In caudal sections, strong PCDH19 expression established a clear dorsal–ventral boundary relative to the ventral PCDH10/PCDH17-dominant territory (Figure 9E,F). The PCDH10/PCDH19 merge highlighted medial–lateral complementarity (Figure 9E), whereas the PCDH17/PCDH19 merge emphasized ventral–dorsal complementarity (Figure 9F). The triple-color merged map confirmed largely complementary territories, with partial overlaps confined to transitional zones (Figure 9G).

## DISCUSSION

Our results indicate that δ2-PCDH-defined molecular domains span the BG, including the striatum, pallidum, and SNr, creating segregated but partially overlapping territories. The graded and complementary expression of PCDH10, PCDH17, and PCDH19 establishes distinct anatomical compartments, providing a potential molecular basis for the organization of parallel BG circuits.

### Molecular Codes Defining Parallel BG Circuits

The δ2-PCDHs define functional territories in the primate BG that are both distinctly and continuously connected, which correspond probably to the limbic, associative, and sensorimotor circuits: PCDH17 is enriched rostrally, likely related to the associative circuit; PCDH10 is predominant in the central, ventromedial domain, likely aligning with the limbic circuit; and PCDH19 is concentrated caudally and dorsolaterally, likely organizing the sensorimotor circuit (Figure 10A). The graded nature of these expression patterns, supported by their complementary distribution, may facilitate smooth transitions and the integration of the circuits. Collectively, the δ2-PCDH molecular topography clearly delineates the historically elusive boundaries of the parallel BG circuits.

**Figure 10.**
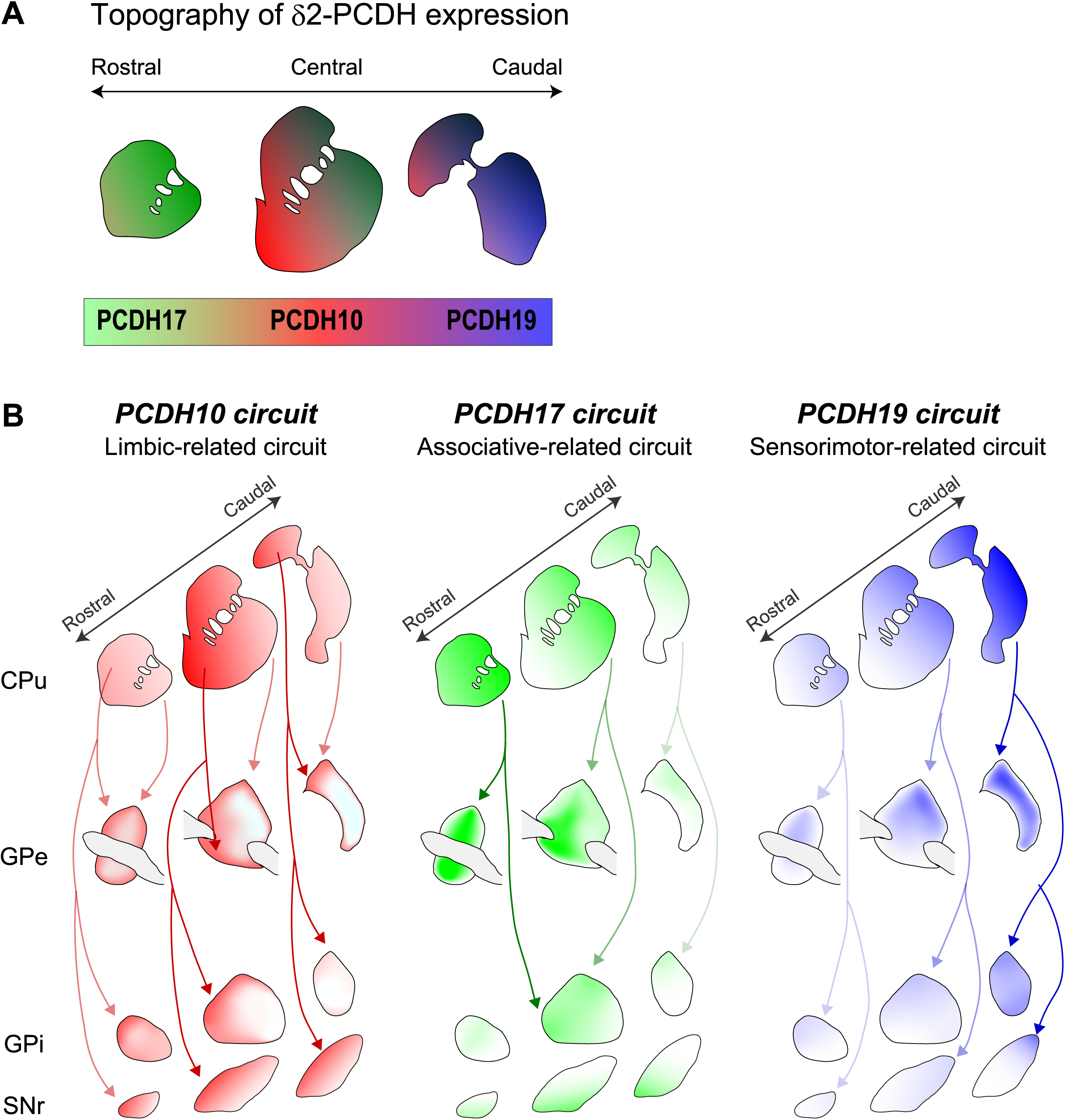
δ2-PCDH molecular code in primate parallel BG circuits. (B) Complementary, region-specific expression patterns of PCDH10, PCDH17, and PCDH19 across the striatum, which broadly correlate with the limbic, associative, and sensorimotor territories, respectively (compare to Figure 1). (C) A molecular wiring diagram illustrating the organization of parallel BG circuits. This diagram is based on a model of δ2-PCDH-mediated selective interactions among the striatum, GPe, and GPi/SNr, proposing molecularly defined PCDH10, PCDH17, and PCDH19 parallel circuits. The arrow intensity represents the strength of individual projections.

The δ2-PCDH territories appear to link striatal inputs to downstream pallidal and nigral outputs in a topographic manner. This is in register with our previous findings on the mouse BG, demonstrating that striatal neurons selectively connect to output nuclei expressing the same δ2-PCDH based on highly specific and homophilic interactions at synaptic sites.^33^ We, therefore, infer that this molecular wiring rule is evolutionarily conserved across species. Following this rule, *PCDH10 circuits* originate in the ventromedial CPu, projecting to the peripheral GPe/VP, rostral GPi, and medial/dorsal SNr (Figure 10B1); *PCDH17 circuits* arise from the rostral CPu, projecting to the rostral inner GPe/VP and ventral SNr (Figure 10B2); and *PCDH19 circuits* emerge from the caudal CPu, projecting to the caudal inner GPe, caudal GPi, and dorsolateral SNr (Figure 10B3). Consequently, δ2-PCDHs likely serve as molecular codes that organize and link striatal inputs to the output nuclei of the BG in a continuous but functionally specialized manner, thereby defining the limbic, associative, and sensorimotor BG circuits.

This topographic molecular architecture may be further subdivided into finer functional territories. In rodents, members of the δ1-PCDH family also exhibit distinct topographic expression patterns that differ from those of δ2-PCDHs.^32^ Therefore, the combination of δ1- and δ2-PCDH expression patterns—that is, the combinatorial expression of non-clustered PCDHs—could provide an additional molecular framework for defining finer functional compartments within the BG. The combinatorial adhesive properties of δ2-PCDHs, together with those of δ1-PCDHs, mediate fine-tuned binding specificity^36,40^ and are likely to specify detailed circuit architectures. Future integrative analyses will be essential to determine whether similar coding principles also operate in the cerebral cortex and thalamus, potentially establishing a unified molecular framework that accounts for the parallel and distributed loop networks of the BG.

### δ-Protocadherins and BG Evolution

The BG are among the most evolutionarily conserved neural systems, preserving their core input–output organization across vertebrates for over half a billion years, from lampreys to mammals.^41–43^ In amniotes (reptiles, birds, and mammals), the striatum became more densely populated and more extensively innervated by cortical projections, governing increasingly complex behaviors.^41–43^ Interestingly, non-clustered PCDHs, including δ1-PCDHs and δ2-PCDHs, form an ancient and conserved family present from lampreys to mammals,^44^ suggesting a preserved molecular function in BG evolution. By contrast, clustered PCDHs emerged later, in the ancestor of jawed vertebrates (Gnathostomes), and were absent in lampreys.^44^ This later emergence likely facilitated such fine circuit tuning through combinatorial diversity, as compared to non-clustered PCDHs.

Parallel BG circuits—limbic, associative, and sensorimotor—are well-established in mammals.^10,12,45^ In rodents, PCDH10 and PCDH17 exhibit complementary expression patterns that effectively segregate these circuits,^33^ while PCDH19 appears in the caudal striatum at relatively low levels.^32^ In primates, the distribution of homologous PCDH10, PCDH17, and PCDH19 is largely conserved. Compared with rodents, however, PCDH19 appears to be more broadly distributed in the sensorimotor territory, a domain occupied more extensively by PCDH10 in rodents. In addition, these δ2-PCDHs form broader and more continuous gradients across the striatum, pallidum, and SNr, reflecting an evolutionary expansion and integration of limbic, associative, and sensorimotor territories. The δ2-PCDH–mediated molecular patterning thus provides a conserved yet flexible system that preserves the core BG architecture while achieving diversification into circuits enabling complex, adaptive behaviors.

### Functional Implications for δ2-PCDH-defined Parallel BG Circuits

The complementary and continuous δ2-PCDH compartments in the BG provide a molecular framework for the functional specialization of the limbic, associative, and sensorimotor circuits. The limbic circuit, aligned with PCDH10-rich ventral territories, processes emotion, motivation, and the expected value of actions as a core component of the reward system.^46^ Supporting evidence comes from *Pcdh10* knockout mice, which exhibit accentuated anxiety- and fear-related behaviors, implicating this molecule in emotional processing.^39^ The associative circuit, corresponding to the rostral CPu and PCDH17-dominant territories, mediates cognitive control and voluntary, goal-directed actions.^22^ It is particularly engaged during early learning, participating in trial-and-error processes that enhance behavioral accuracy.^47^ Consistent with this, *Pcdh17* knockout mice show deficits in stress coping,^33^ and human *PCDH17* variants are associated with reduced cognitive performance and altered personality traits, suggesting a molecular contribution to cognitive regulation.^48^ Finally, the sensorimotor circuit, encompassing the caudal CPu and PCDH19-dominant regions, mediates habitual and automatic behaviors.^1,21^ Its activity increases during late learning, facilitating rapid and efficient motor execution.^47^ Clinical phenotypes associated with *PCDH19* mutations—including cognitive and neurodevelopmental impairments—are consistent with deficits in sequencing, coordination, and other sensorimotor functions.^49^ Together, the δ2-PCDH system may ensure functional segregation through homophilic neuronal adhesion and synaptic regulation within each molecular territory, while enabling the BG to coordinate the sequential activation and integration of these parallel circuits.

### Neurological and Psychiatric Implications for δ2-PCDH-defined Parallel BG Circuits

The δ2-PCDH-defined parallel BG circuits provide crucial insights into neurological and psychiatric disorders associated with BG dysfunctions. These conditions are now understood not as uniform impairments of a single circuit, but as disruptions across multiple parallel circuits.^11^ Within this framework, individual symptoms—even when shared across different diagnoses—can be mapped onto the topographically-organized limbic, associative, and sensorimotor circuits. In Parkinson’s disease, motor deficits are derived from dysfunctions of the sensorimotor circuit, whereas non-motor symptoms such as depression and impaired decision-making involve the limbic and associative circuits.^16,50^ In ASD, social communication deficits are linked to the limbic circuit, cognitive inflexibility to the associative circuit, and stereotyped behaviors and sensory hypersensitivity to the sensorimotor circuit.^51^ In OCD, excessive fear and anxiety reflect hyperactivity of the limbic circuit, repetitive thoughts and impaired cognitive control involve the overactive associative circuit, and compulsive behaviors depend on the sensorimotor circuit.^9^ Similarly, in schizophrenia, cognitive deficits primarily stem from associative circuit dysfunctions, while affective symptoms involve the limbic circuit.^10^

These circuit-specific impairments may be associated with molecular pathologies involving abnormalities in δ2-PCDHs (PCDH10, PCDH17, and PCDH19), which can disrupt selective connectivity and functional integration within the BG networks. Indeed, in humans, *PCDH17* mutations have been associated with schizophrenia and mood disorders;^52,48^ *PCDH10* mutations with ASD and OCD;^53–55^ and *PCDH19* mutations with epilepsy, cognitive impairments, and neurodevelopmental abnormalities.^49,56^ The δ2-PCDH–based molecular framework of parallel BG circuit organization thus provides a unifying link between the molecular pathology and the circuit-specific symptomatology observed across neurological and psychiatric disorders. Moving forward, integrative analyses across molecular, cellular, and systems levels will be essential to elucidate the underlying pathophysiology and to develop targeted and circuit-based therapeutic strategies.

### Limitations of the study

A primary limitation of this study is the exclusive use of infant macaques, which prevents direct confirmation of whether the observed complementary δ2-PCDH expression patterns persist into adulthood. Nevertheless, the use of infant subjects was essential as δ2-PCDH expression peaks during early synaptogenesis. Furthermore, we hypothesize that the fundamental parallel organization of basal ganglia circuits is established prior to the maturation of higher cognitive functions, making infant subjects appropriate and sufficient for verifying this basic circuit architecture. Future research is required to track the long-term maintenance of this molecular code in mature primates.

## RESOURCE AVAILABILITY

### Lead contact

Further information and requests for resources should be directed to and will be fulfilled by the lead contact, Naosuke Hoshina.

### Materials availability

The primary antibodies used in this study are available from the lead contact upon reasonable request. No new unique reagents were generated in this study.

### Data and code availability

No new code was generated in this study. Any additional information required to reanalyze the data reported in this paper is available from the lead contact upon request.

## ACKNOWLEDGMENTS

The authors would like to thank Ken-Ichi Inoue, Kiyomi Nagaya, Teiko Kuroda, Aja Sanzone for their assistance in the data collection. This work is supported by the JSPS KAKENHI Grant Number 26430046 (N.H.), 24K02133 (N.H.), the Uehara Memorial Foundation (N.H.), the Brain Science Foundation (N.H.), the Naito Foundation (N.H.), the Astellas Foundation for Research on Metabolic Disorders (N.H.), Takeda Science Foundation (N.H.), and internal funding from the Okinawa Institute of Science and Technology Graduate University, Japan (T.Y.).

## AUTHOR CONTRIBUTIONS

Conceptualization, N.H. and M.T.; methodology, N.H. and M.T.; investigation, N.H., M.H., T.Y. and M.T.; writing – original draft, N.H.; writing – review and editing, N.H. M.H., and M.T.; funding acquisition, N.H. and T.Y.; supervision, N.H. and M.T.

## STAR★METHODS

### KEY RESOURCES TABLE

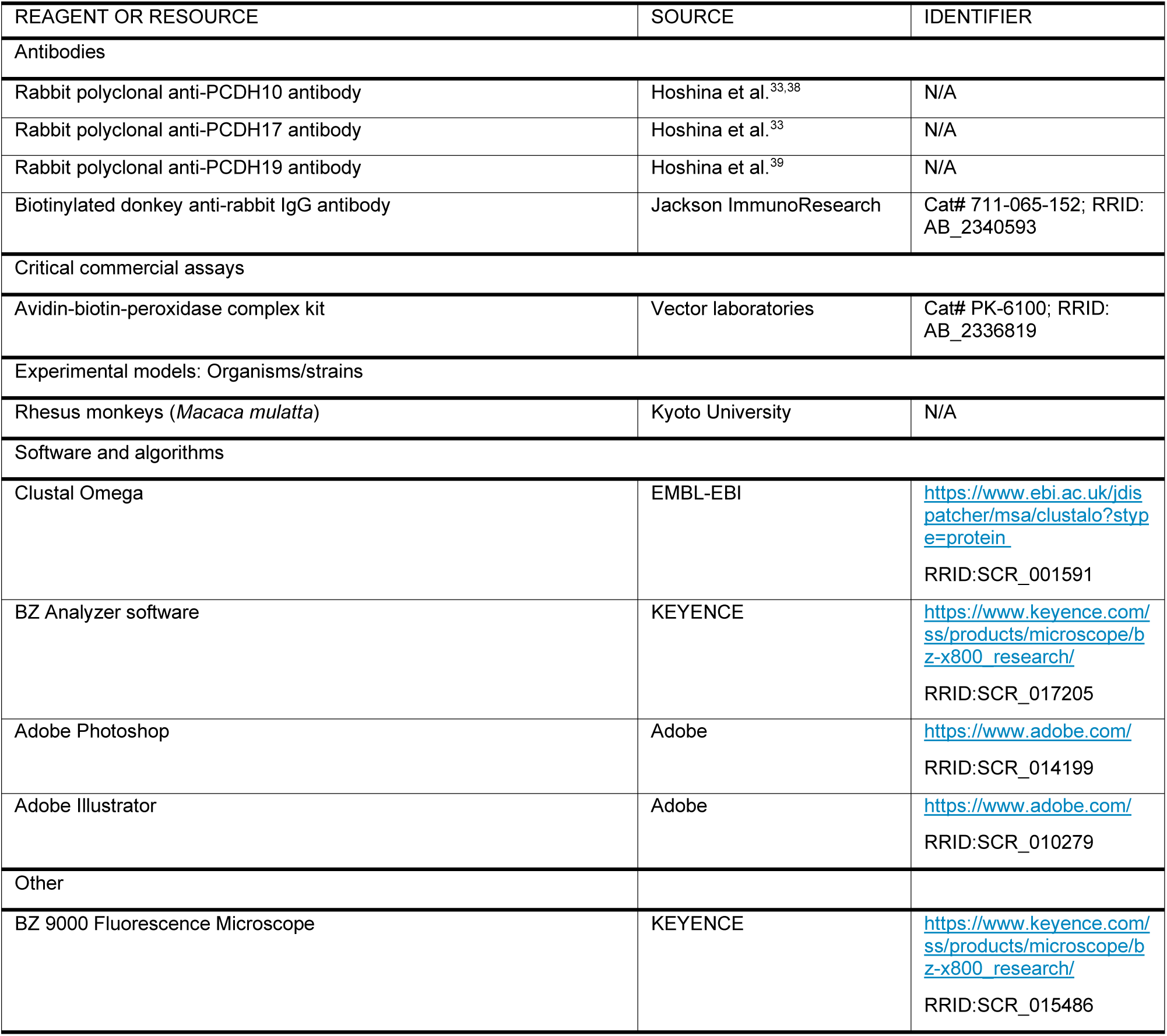

### EXPERIMENTAL MODEL AND STUDY PARTICIPANT DETAILS

#### Animals

Infant rhesus monkeys (*Macaca mulatta*) of either sex, aged 1–2 months, were included in the study. All experimental procedures were approved by the Institutional Animal Care and Use Committee of the Center for the Evolutionary Origins of Human Behavior, Kyoto University, and were conducted in accordance with the Guidelines for the Care and Use of Nonhuman Primates (Center for the Evolutionary Origins of Human Behavior, Kyoto University).

### METHOD DETAILS

#### Antibodies

Rabbit polyclonal antibodies against PCDH10, PCDH17, and PCDH19 were custom generated in our laboratory and have been extensively characterized in previous studies.^33,38,39^ The specificity of each antibody was validated by the complete absence of staining in the corresponding knockout mice, which served as negative controls. For immunohistochemistry, antibodies were used at concentrations of 1–4 µg/ml.

#### Immunohistochemistry

Monkeys were initially sedated with ketamine hydrochloride (10 mg/kg, i.m.) and xylazine hydrochloride (1–2 mg/kg, i.m.), followed by deep anesthesia with an overdose of sodium pentobarbital (50 mg/kg, i.v.) for transcardial perfusion. Animals were perfused with 0.1 M phosphate-buffered saline (PBS, pH 7.4), followed by 10% formalin in 0.1 M phosphate buffer (PB, pH 7.4). Brains were removed from the skull, post-fixed in the same fixative (fresh) overnight at 4 °C and saturated with 30% sucrose at 4 °C.

Coronal sections (60 μm) were cut serially on a freezing microtome. Every eighth section was processed for immunohistochemical staining using the standard avidin-biotin-peroxidase complex (ABC) method. Sections were blocked in 1% skim milk and incubated for three days at 4 °C with primary antibodies (PCDH10, PCDH17, or PCDH19) in PBS containing 0.1% Triton X-100 and 2% normal donkey serum. Following primary incubation, sections were treated with biotinylated secondary antibodies (Jackson ImmunoResearch) and subsequently incubated with ABC Elite solution (Vector Laboratories). Antigen visualization was performed using 50 mM Tris-HCl (pH 7.6) containing 0.04% diaminobenzidine, 0.04% nickel chloride, and 0.002% hydrogen peroxide.

#### Image analysis

Sections were imaged using a BZ-9000 All-in-One Fluorescence Microscope (Keyence). All images were obtained under identical acquisition settings, including exposure time, detector gain, and offset, to ensure comparability. Wide-field images were acquired by stitching several adjacent images together using the BZ Analyzer Software (Keyence). For visualization, the contrast of single-stained images was adjusted using Adobe Photoshop (Adobe) so that the signal intensity in the corpus callosum was uniform across all sections, serving as a background reference. The single-stained images were then color-assigned and merged to visualize the distinct expression patterns of PCDH10, PCDH17, and PCDH19 across BG regions, including the striatum and output nuclei. Color-coded merged images were generated in Adobe Illustrator (Adobe) using the “Multiply” blend mode function. All analyses presented were qualitative, and no quantitative measurements were conducted.

